# RLIM enhances BMP signalling mediated fetal lung development in mice

**DOI:** 10.1101/507921

**Authors:** Molka Kammoun, Elke Maas, Nathan Criem, Joost Gribnau, An Zwijsen, Joris Robert Vermeesch

## Abstract

RLIM, also called RNF12, is an X-linked E3 ubiquitin ligase. It is a pleotropic protein acting as a negative regulator of LIM-homeodomain transcription factors and of SMAD7, an antagonist of the BMP and TGFβ pathways. Mutations in *RLIM* are associated with variable congenital malformations in human. The molecular mechanism causing these malformations is still poorly understood.

In this study, we used a conventional knockout (KO) mouse model to phenotypically characterize male mutants and to investigate the effect of *Rlim* mutation on BMP pathway signalling. Only 50% of KO male neonates survive and are significantly smaller. We show that *Rlim* deletion alters lung branching and maturation resulting in severe respiratory distress and neonatal death. Except for a single case of cardiac interventricular septal defect, no other malformations that mimic the human condition were observed. Western blot analysis shows that RLIM deficiency is associated with an increase of its target SMAD7 levels and a loss of SMAD1/5/9 activation. Interestingly, SMAD5 levels were diminished, suggesting another regulatory loop between RLIM and the BMP pathway.

In conclusion, RLIM is important for lung development in mice and exerts this function in part through intracellular propagation of BMP signalling.

**Author summary:** *RLIM* is an X-linked gene encoding an enzyme that regulates a number of proteins including SMAD7, an antagonist of BMP pathway. *RLIM* mutations are associated with intellectual disability and several congenital malformations in boys. In mice, RLIM is thought to be nonessential for embryos development. Using a mouse model in which *Rlim* is deleted, we show that 50% of males carrying the deletion display respiratory distress and die short after birth. No congenital malformations mimicking the human phenotype have been observed. Histological analysis of lung samples show that *Rlim* is critical for late lung development and maturation. The analysis of lung samples by western blot, a technique that quantifies proteins, is consistent with a dysregulation of BMP pathway. These finding confirm the importance of RLIM for murine embryos development and provide a link between human *RLIM* pathology and BMP pathway.

## Introduction

*RLIM*, also called *RNF12*, is an X-linked gene that encodes a widely expressed RING-H2 zinc finger protein acting as an E3 ubiquitin ligase [1]. RLIM was first discovered as a LIM homeodomain transcription factor repressor. Later studies have demonstrated a wide range of RLIM functions including effects in cancer. For instance RLIM plays a role in telomere length homeostasis through negative regulation of the telomeric protein TRF1 [2], promotes p53 dependent cell growth and proliferation suppression [3] and enhances TGFβ signal transduction in human osteosarcoma cell lines, thereby activating TGFβ induced migration of these cells [4]. The latter function is achieved through interaction with Smad-ubiquitin regulatory factor 2 (SMURF2), a ubiquitin ligase that inhibits the TGFβ pathway [4]. Furthermore, RLIM enhances the TGFβ and BMP signaling pathways in mouse embryonic stem cells and zebrafish embryos by targeting SMAD7, an inhibitory SMAD, for degradation [5]. Moreover, RLIM is a sex-specific epigenetic regulator of nurturing tissues in pregnant and lactating female mice. Whilst the *Rlim* paternal allele is important for mammary gland alveolar cells survival and activity [6], the maternal allele is critical for trophoblast development in female embryos through imprinted X chromosome inactivation (XCI) regulation [7]. Studies on *in vitro* differentiated embryonic stem cells have shown that RLIM stimulates random XCI through degradation of REX1 that inhibits Xist (X inactivation specific transcript) transcription [8]. Very recently, Gontan and collaborators have shown that RLIM mediated REX1 degradation is a critical event in imprinted XCI, as the lethal phenotype of Rnf12 ^-/+^ and Rnf12^-/-^ females is rescued in a mutant *Rex1* background [9]. Of interest a maternally transmitted *Rlim* deletion causes not only a complete loss of female carriers because of defective imprinted XCI, but also about half of their male KO pups [9]. Similar results have been observed in an earlier study using conditional *Rlim* KO in oocytes [7]. The male lethality was not the main focus of these studies and was not investigated. However, Shin and collaborators have dissected embryos of crosses between *Rlim* heterozygous females and WT males, from preimplantation stage up to E10.5 stage, and have not reported a significant anomaly in male embryos [7] suggesting a late developmental defect. By contrast, conditional *Rlim* KO in epiblast and in its derived tissues does not lead to lethality in males or in females [10]. This finding indicates that RLIM would only be important for imprinted XCI [10] and suggests that KO male mortality is due to an RLIM deficiency in the non-epiblast tissues, the trophectoderm (trophoblast and extraembryonic ectoderm) and/or primitive endoderm (parietal and visceral endoderm) derived lineages. Interestingly, the visceral endoderm does not only give rise to extraembryonic tissues, but also contributes to definitive endoderm in the embryo proper [11].

Contrary to mouse models, human *RLIM* mutations have been shown to cause a large phenotypic spectrum in male patients. Mutations in *RLIM* have first been reported to cause X-linked intellectual disability and behavioral disorders [12,13]. Most recently, we have provided a detailed phenotypic description of 9 families with rare and likely pathogenic *RLIM* variants [14]. Next to the aforementioned phenotype, *RLIM* lesions are also associated with multiple congenital malformations which are dominated by congenital diaphragmatic hernia and lung hypoplasia, in addition to variable congenital heart defects, urogenital tract and skeletal abnormalities. These malformations are compatible with a dysregulation of BMP dependent signaling which is normally promoted by RLIM-mediated degradation of SMAD7 [5,14].

BMPs are ligands of the transforming growth factor β (TGFβ) family, which also includes TGF-β, growth and differentiation factors, activins and nodal. The binding of BMPs to heteromeric type I and type II transmembrane serine/threonine kinase receptors, triggers activation by phosphorylation of the so-called receptor-regulated SMADs (R-SMADs), SMAD1, SMAD5 and SMAD9 (formerly named SMAD8). Phosphorylated R-SMADs form a complex with SMAD4 and translocate into the nucleus to modulate target genes expression. Inhibitory SMADs, SMAD6 and SMAD7, antagonize amongst others R-SMADs phosphorylation [15].

Building further upon the human mutant *RLIM* phenotypic spectrum and on the results of the previously described *Rlim* mouse models, we hypothesize that lethality of male mice results from severe malformations in endodermal tissues such as liver and digestive tract that contain visceral endoderm derivatives; or in tissues relying on visceral endoderm signaling in the early embryo such as the forebrain, the heart and the septum transversum, one of the embryonic sources of the diaphragm [16,17]. The objective of this study is to better understand RLIM function in mammalian organogenesis and its molecular mechanism. To this end, we used a conventional *Rlim* KO mouse model to phenotypically characterize male carriers and to investigate the BMP pathway in affected tissues.

## Results

### *Rlim* deletion causes a growth delay and neonatal lethality

To determine the distribution of the *Rlim* (*Rnfl2*) deletion, 19 crosses of *Rnf12*^-/y^ KO males and *Rnf12*^+/+^ WT females (in the -/+ or +/- nomenclature, the maternally inherited allele is shown first), and 25 crosses of heterozygous *Rnf12*^+/-^ females and *Rnf12*^+/y^ WT males were performed. All P1 pups born to *Rnf12*^-/y^ KO males and WT females showed normal sex ratios and a Mendelian distribution of the offspring (63 *Rnf12*^+/-^ females, 61 *Rnf12^+/y^* males). Conversely, a maternally transmitted *Rnf12* KO allele leads to a complete absence of mutant female pups *Rnf12*^-/+^ and to a skewed male (*Rnf12^+/y^*/*Rnf12*^-/y^) ratio (56 WT, 27 KO p= 0.001) (Fig 1). In agreement with the previous studies, surviving *Rnf12*^-/y^ males were all fertile (n=27). The dissection of ten survivors did not reveal gross congenital malformations. Furthermore, the number of live pups on the day of birth and at weaning were comparable, suggesting a prenatal or a neonatal lethality of half of the *Rnf12* KO males.

**Fig 1:**
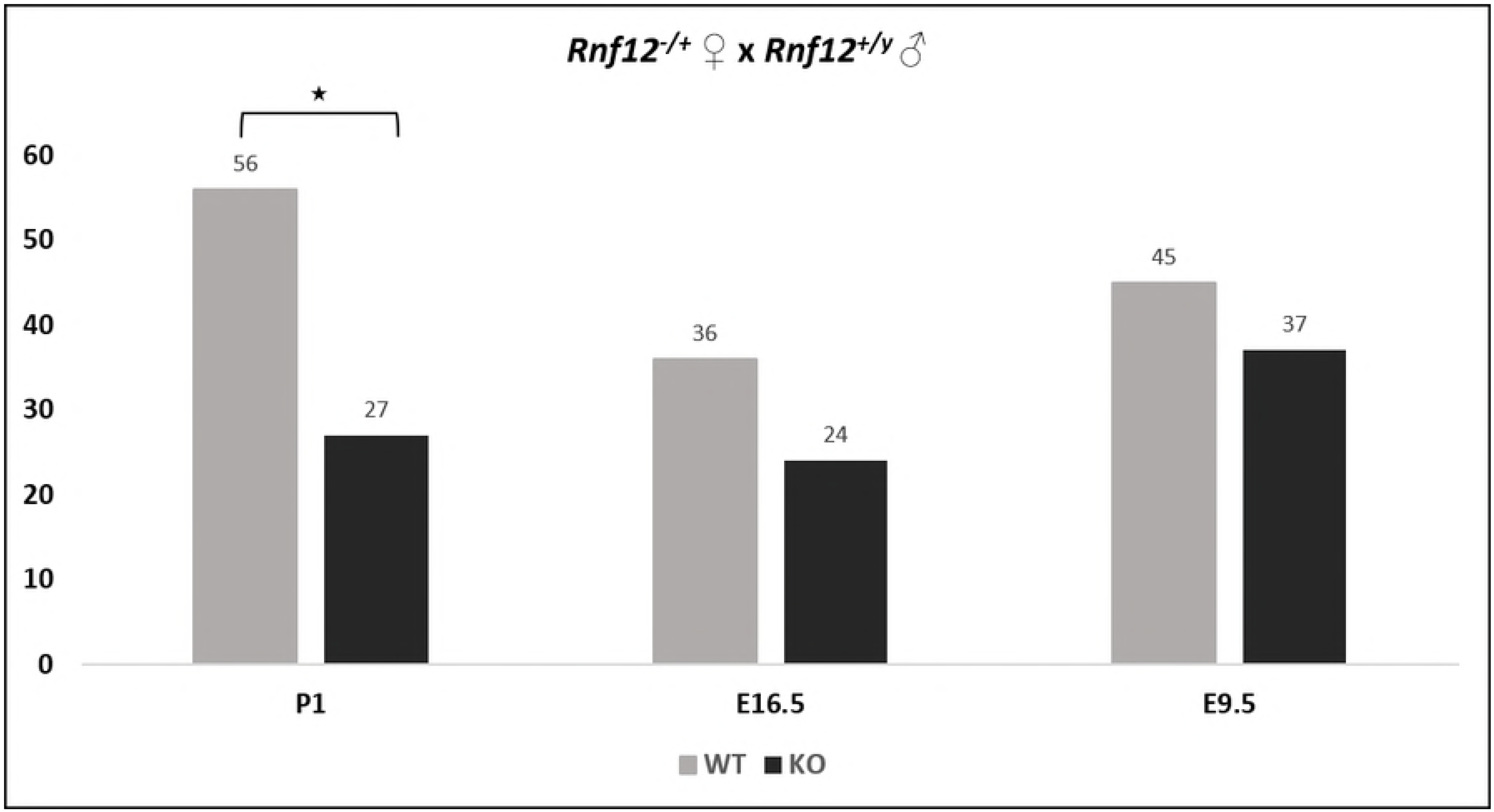
A maternally transmitted *Rlim* deletion leads to a perinatal lethality in males. Schematic diagram for male pups and embryos (E9.5 and E16.5) generated by Rnf12^-/+^ female and WT male crosses. *:0.05<p<0.001

To better define the lethality stage, we dissected embryos at E9.5 and at E16.5. All but two *Rnf12*^-/+^ females died before E16.5. The female phenotype will not be discussed in this paper. As expected for males, numbers of WT *Rnf12^+/y^* and *Rnf12*^-/y^ KO male E9.5 embryos were not significantly different (45 WT, 37 KO) and their external examination did not reveal specific anomalies in *Rnf12*^-/y^ embryos. Likewise, E16.5 *Rnf12*^-/y^ were indistinguishable from WT male feti (36 WT, 24 KO). Despite a broad *Rlim* expression in many different tissues [10], microdissection of seventeen E16.5 KO males excluded gross malformations. Specifically, diaphragm, lung lobulation, anatomical heart orientation, liver size and lobulation, digestive tract, kidneys and adrenal glands looked normal in all the feti. These results suggest that for this mouse model, KO male die perinatally.

We therefore proceeded to dissections at E18.5. Following cesarian section, a total of 56 feti were monitored for 1 hour and were weighed. All the pups were viable as indicated by a beating heart at the extraction moment. Surprisingly, all but one *Rnf12*^-/y^ pup (16/17) showed respiratory failure signs: cyanosis, gasps with costal retraction, and deceased within 30 minutes (n=16). KO male lungs had a normal size and lobulation, but looked paler. Contrary to WT lungs, KO male lungs sank in PBS confirming the absence of air inflation (Fig 2). In addition, KO males were significantly smaller than WT feti (p< 0,0001). Figure 3 shows fetal weight distribution across the monitored in-utero euthanized feti (n= 17 *Rnf12*^-/y^ and 25 WT males).

**Fig 2:**
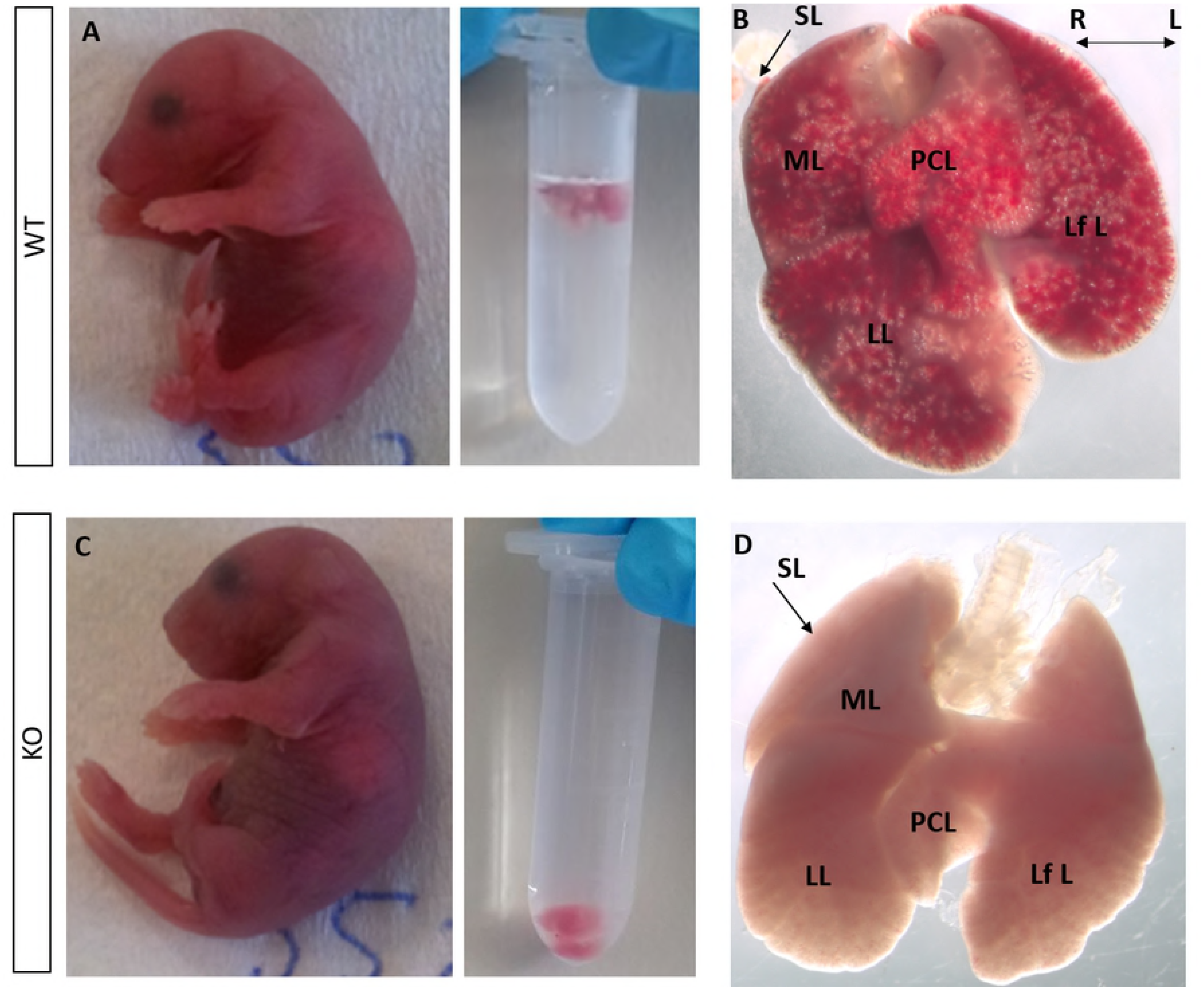
*Rnf12^-/y^* KO males display sings of respiratory distress. Representative pictures of WT and *Rnf12^-/y^* KO E18.5 male feti and corresponding lung float test (A and C). Note that KO fetus is cyanotic with a negative float test in PBS confirming the absence of alveoli inflation. B and D: representative pictures of WT and KO lungs. Note the normal size and lobulation of KO lungs. Abbreviations: SL: superior lobe, ML: middle lobe, LL: lower lobe, PCL: post-caval lobe, LfL: left lobe

**Fig 3:**
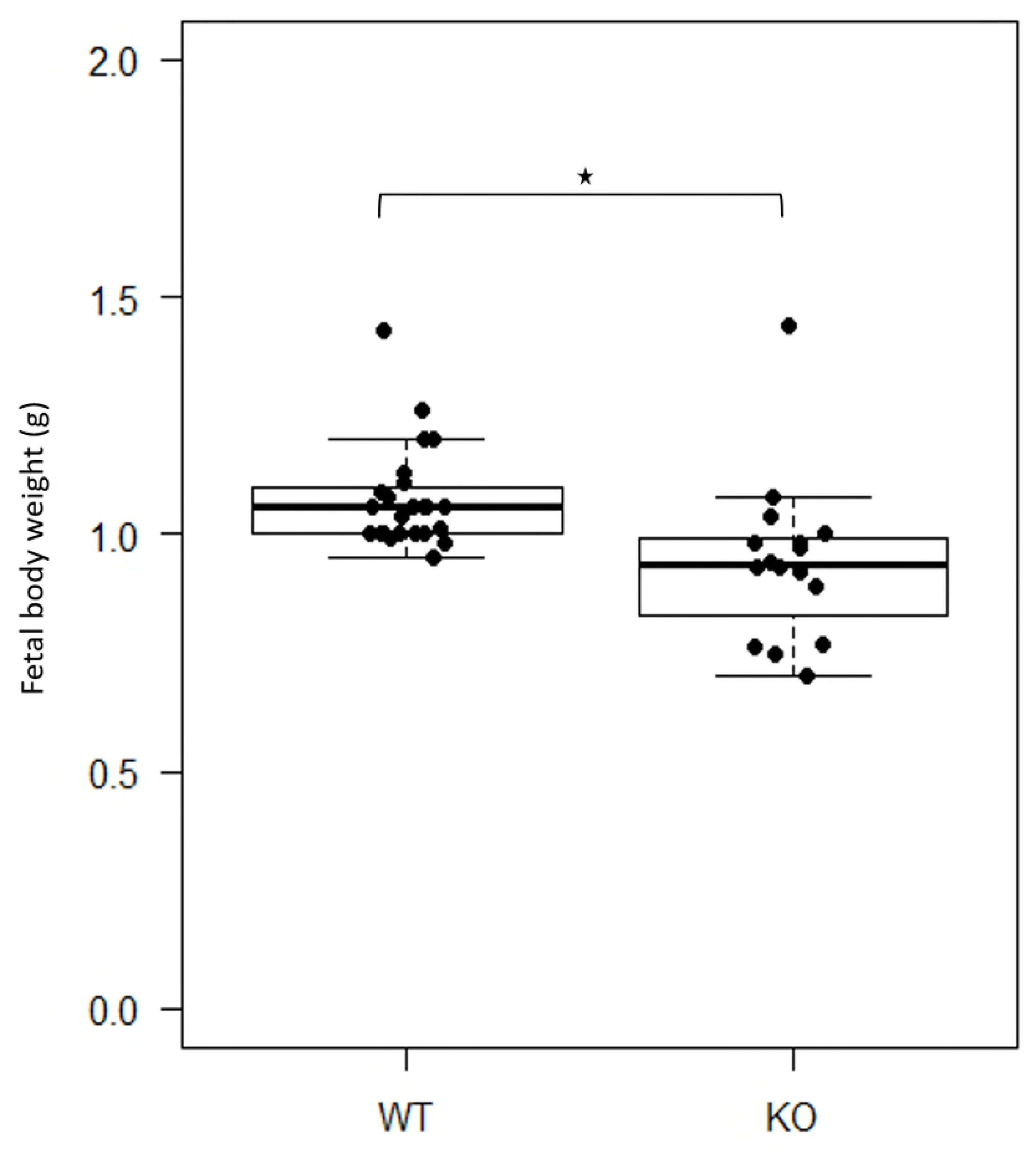
*Rnf12^-/y^* KO males have a smaller birth weight. Fetal body weight distribution across the monitored in-utero euthanized feti (n= 24 WT and 17 *Rnf12^-/y^* KO males)

### Deletion of *Rlim* causes fetal lung branching and maturation anomalies and heart septation defect

Four *Rnf12*^-/y^ KO and five WT males from 4 different E16.5 litters were randomly selected and tissues were subjected to histological examination. The liver and the diaphragm were comparable and normal in both groups (Fig 4). A ventricular septal defect was identified in one *Rnf12*^-/y^ fetus (Fig 4). Wild type fetal lungs showed a pseudoglandular structure. Developing airways were more dilated in two *Rnf12*^-/y^ KOs which is consistent with a delayed branching (Figs 5A-B).

**Fig 4:**
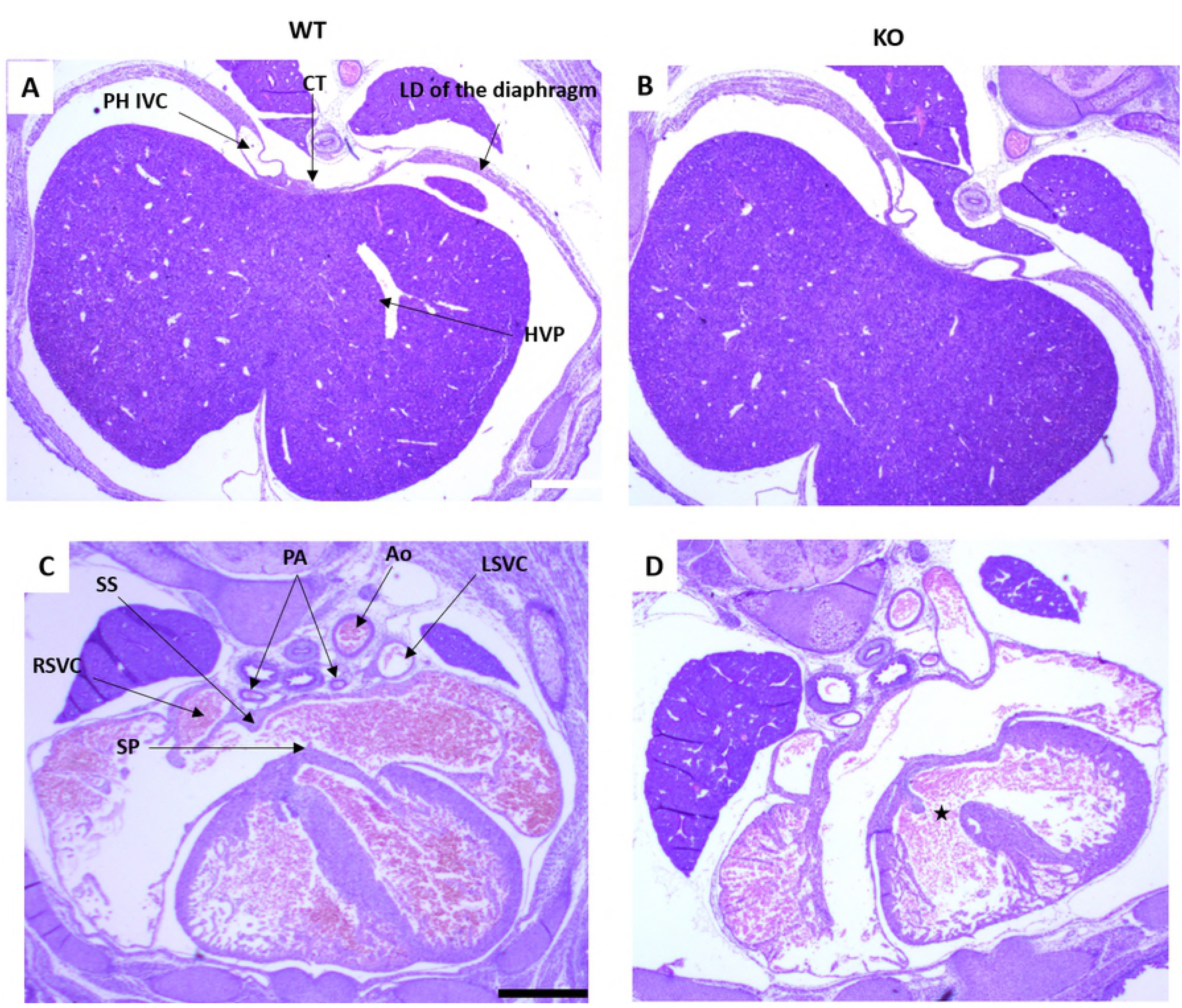
Haematoxylin-eosin staining of a transverse section through the liver and the heart of E16.5 feti: (A) and (C): *Rnf12^+/y^* WT. (B) and (D): *Rnf12^-/y^* KO (Scale bar= 500 μm). A ventricular septal defect is shown in D by the star. Abbreviation: Ao: aorta, CT: central tendon, HVP: hepatic venous plexus, LD: left dome, LSVC: left superior vena cava, RSVC: right superior vena cava, PH IVC: post hepatic inferior vena cava, PA: pulmonary arteries, SP: septum premium, SS: septum secundum,

**Fig 5:**
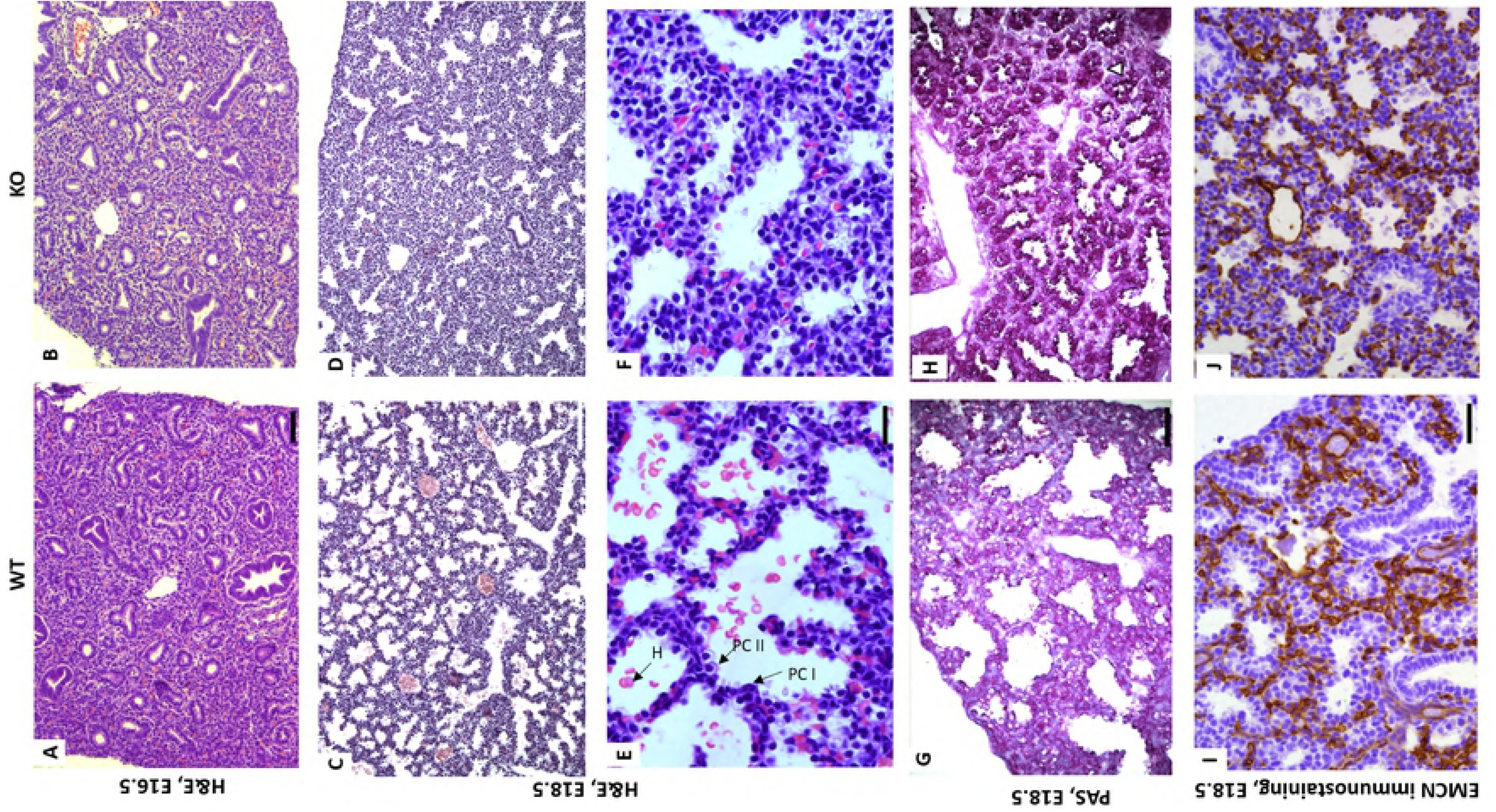
*Rnfl2* deletion causes a delayed lung branching and maturation and an abnormal epithelial differentiation which is not due to impaired vascular development. (B) Reduced lung branching in *Rnf12^-/y^* KO feti is shown by enlarged airways at E16.5 (scale bar=100 μm). (D) Reduced terminal saccules size and number at E18.5 (scale bar=100 μm). (H) increase of PAS staining in KO lung tissue as shown by the black arrow (scale bar= 100 μm) reflects a delayed surfactant production. (E) (F) Pneumocytes type I (PC I) and type II (PC II) are shown. Pneumocytes II are more abundant and larger in *Rnf12^-/y^* KO feti indicating an abnormal epithelial differentiation (scale bar=20 μm). The abnormal presence of hematocytes is observed in both groups. (I) (J) Endomucin immunostaining marking endothelial cells did not show a difference between *Rnf12* WT and KO feti (scale bar= 100 μm). *: 0.05> p> 0.001.

Lung development was further investigated at E18.5, when the lungs are at the saccular stage (Fig 5C). To examine prenatal lung morphology before air exposure, E18.5 feti were euthanized *in utero* [18]. Standard histologic analysis was performed on lung samples collected from 8 *Rnf12*^-/y^ KO and 8 WT feti from 3 different litters. The distal part of the lungs of 7/8 *Rnf12*^-/y^ feti had a reduced number of terminal sacs; the mesenchymal tissue was thickened and the interalveolar septae were thicker compared to the WT lungs (Figs 5C-D). This abnormal branching was confirmed by quantification of the overall air space area to the entire tissue area ratio which was significantly reduced in *Rnf12*^-/y^ KO (p= 0,0029) (Fig 6). The epithelium lining the primitive alveoli in WT mice contained squamous pneumocytes type I and more cuboidal pneumocytes type II. In *Rnf12*^-/y^ lungs, cuboidal cells were more abundant and larger in some areas, which supports an impaired epithelial differentiation (Figs 5E-F). The uncommon presence of hematocytes was observed in the mesenchyme of both groups and is likely due to the intrauterine euthanasia that causes a hypoxic condition.

**Fig 6:**
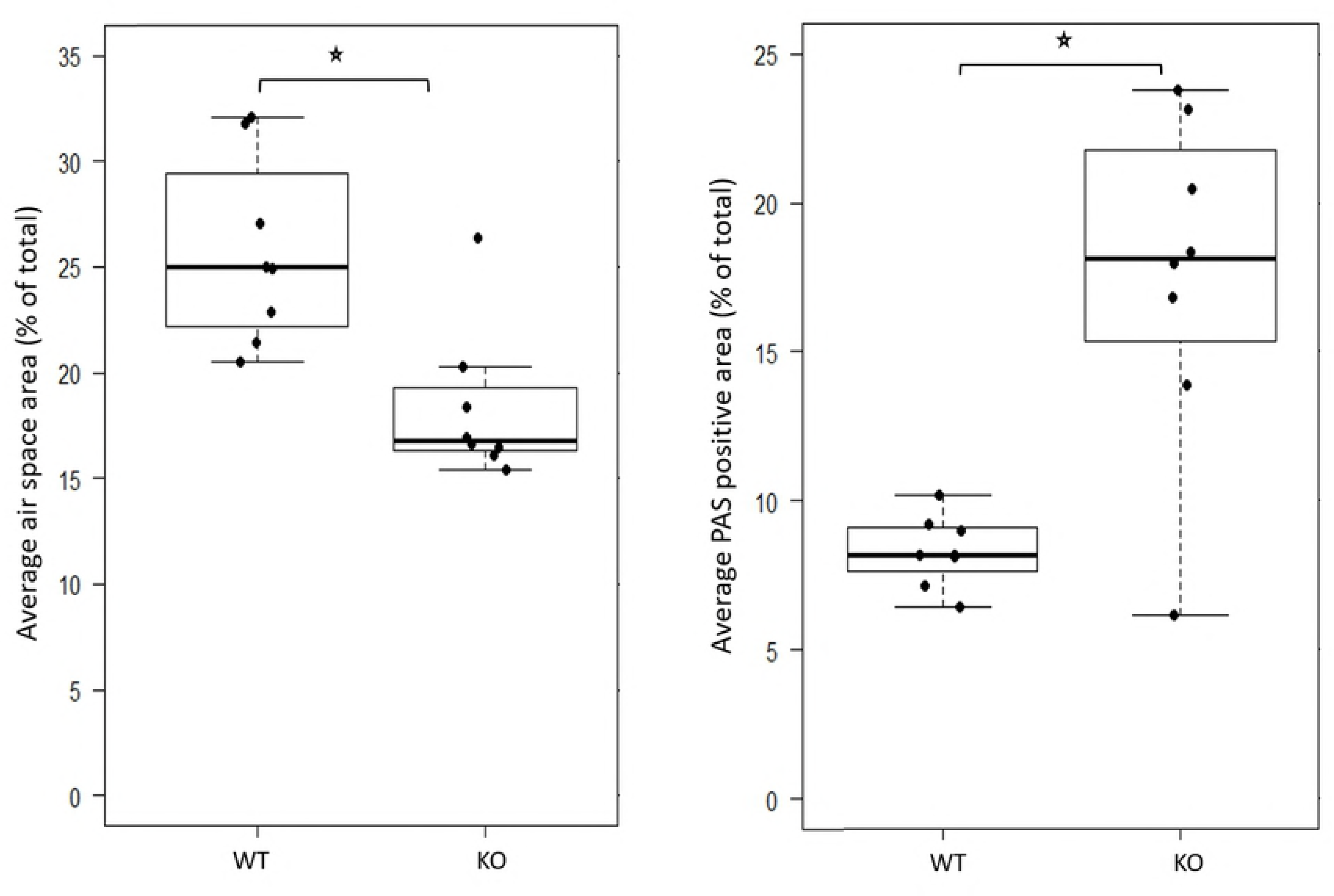
*Rnf12* deletion is associated with a significant reduction of average air space and maturation. Quantification of the average air space and of PAS positive area.

Periodic Acid Shiff staining (PAS) showed a significant increase of surfactant precursor glycogen levels (PAS positive area) in 7/8 *Rnf12*^-/y^ lungs compared to control littermates (p= 0,01), compatible with delayed lung maturation (Fig 5G-H, Fig 6).

Given the paleness of KO lungs, we investigated the pulmonary vascular development using anti-endomucin anti-CD31 antibodies. No significant difference was observed in capillary density between WT and KO (Figs 5I-J) (data not shown for CD31).

### *Rlim* deletion blunts BMP pathway activity in lungs

Considering that RLIM targets SMAD7 for degradation and consequently potentiates the BMP pathway [5], and that BMP signaling is involved in heart and lung development [19,20], we further investigated this pathway by immunoblotting equal amounts of protein extracted from E18.5 lung samples of 6 KO and 6 WT feti from 4 different litters. These feti were monitored prior to lung extraction and their genotype was confirmed by immunoblotting using an anti-RLIM antibody (Fig 7A). We analyzed canonical BMP effector signaling Phosphorylated SMAD1/5/9, the BMP pathway antagonist SMAD7, and the BMP target gene ID3. Phosphorylated SMAD1/5/9 was drastically reduced in KO lung samples (Fig 7A). Remarkably, ID3 was only slightly reduced in *Rnf12*^-/y^ KO lung samples (Figs 7A, 8) suggesting that ID3 is also regulated by non-SMAD mediated signaling. As expected from the absence of RLIM, SMAD7 was increased in KO males, and this increase was found to be statistically different (Figs 7A, 8).

**Fig 7:**
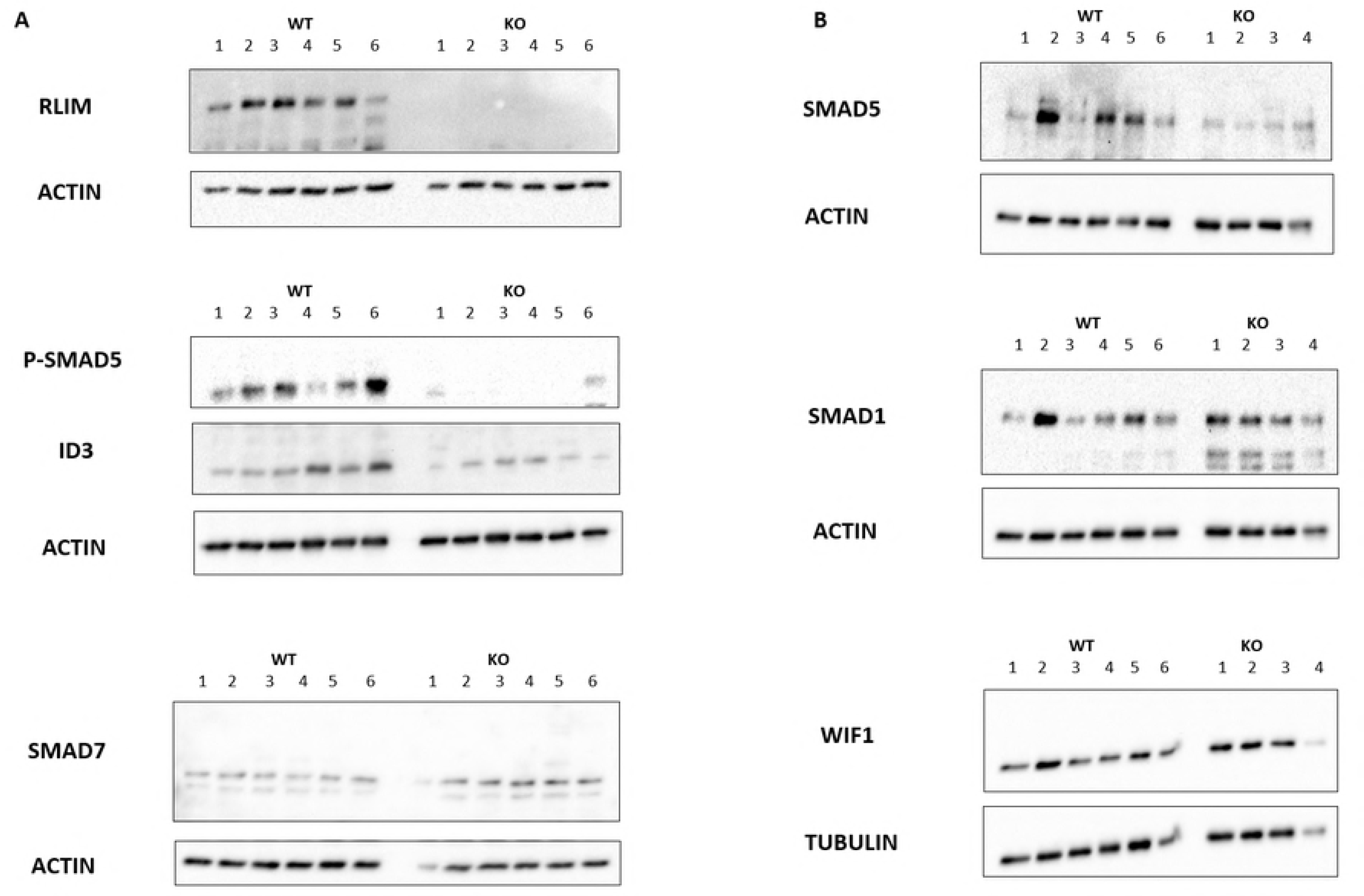
*Rnfl2* deletion attenuates BMP pathway activity. (A) Lung lysates analysis by immunoblots of 6 different WT and *Rnf12^-/y^* KO male feti (E18.5) with the indicated antibodies. (B) Representative pictures of the analysis of 6 KO and 12 WT. The WT and KO lysates were used in the same order in all the experiments.

**Fig 8:**
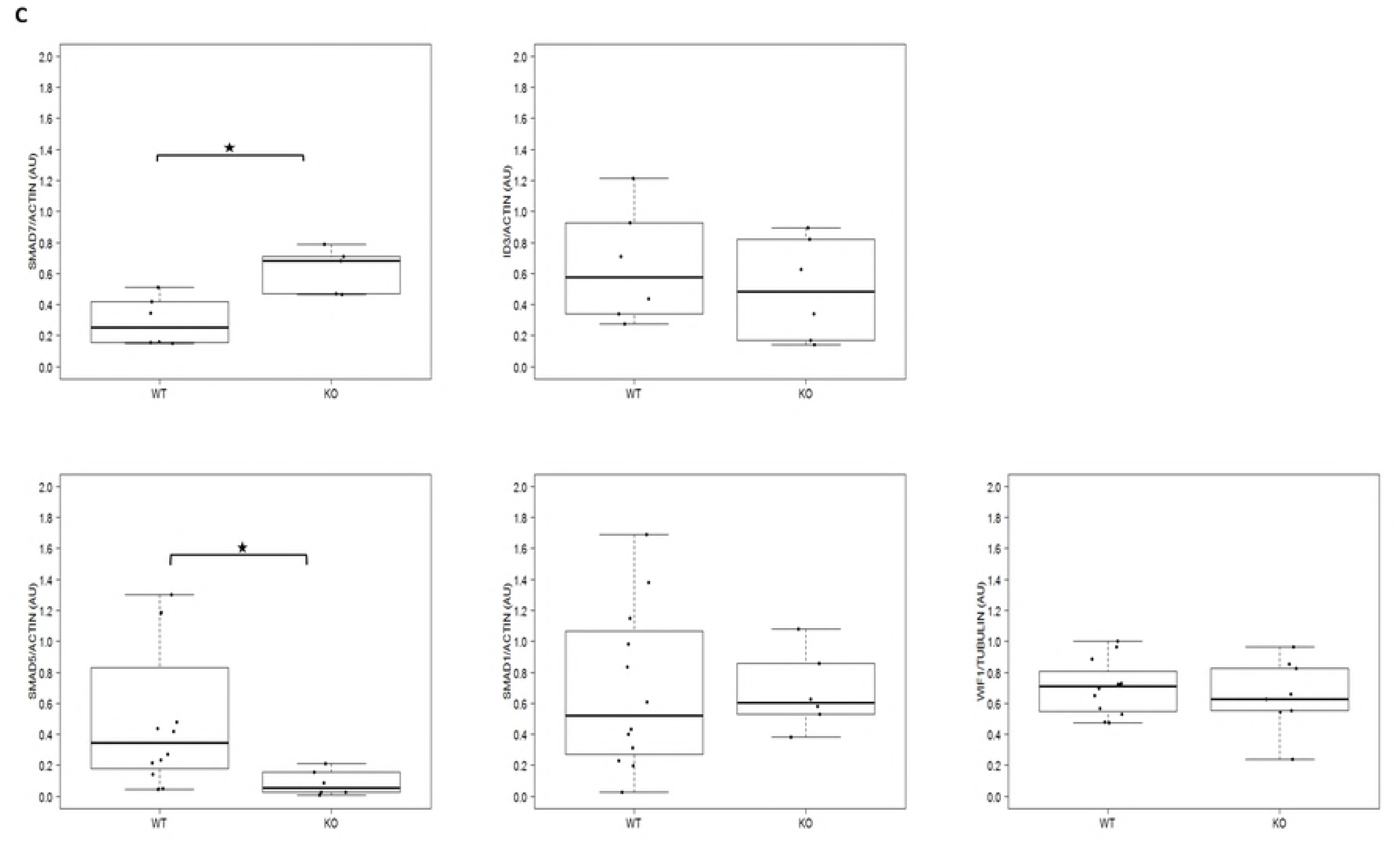
Relative quantification of SMAD7, SMAD5, SMAD1 and WIF1 in two different experiments. *: 0.05> p> 0. 001

Next, to further understand the molecular mechanism underlying this defective BMP signaling and since SMAD1 and its target effector WIF1 (WNT Inhibitory Factor I) are involved in lung maturation [20], we measured the unphosphorylated SMAD1 and SMAD5 and WIF1 levels. Given the variability of these proteins’ level, 6 more WT littermates were analyzed by western blot analysis. Surprisingly, whilst SMAD5 was significantly reduced in KOs, SMAD1 and WIF1 were equally expressed in both groups (Figs 7 B, 8).

## Discussion

Human RLIM is a pleotropic protein and *RLIM* lesions in humans are associated with several congenital malformations including congenital diaphragmatic hernia, various congenital heart defects, ambiguous genitalia, renal and skeletal malformations [14]. These malformations are likely to result from a dysregulated BMP pathway that is involved in several developmental aspects. For instance, BMP4 loss of function leads to syndromic CDH [21]. In addition, BMP4 has a close interaction with *GATA4* which is also critical for diaphragm, lung and for heart development [22]. Urogenital malformations can also be linked to impaired BMP signalling. In fact, kidneys of heterozygous null mutant (*Bmp4*+/-) mice contain multicystic dysplastic regions resulting from an increased apoptotic activity in the metanephric mesenchyme [23]. In a Brazilian population, a number of SNPs in *BMP4* are significantly associated with multicystic dysplastic kidneys disease and ureteroplevic junction obstruction [24]. In addition *BMP* pathogenic variants have been reported association with hypospadias or with 46,XY disorders of sexual development [25,26].

On the other hand, murine RLIM has long been considered to be essential solely for female mouse trophoblast development regulating imprinted XCI. In their recent study, Gontan et al have shown that despite rescue of imprinted XCI in Rnf12:Rex1 double knockout mice, these mice still display loss of male and female offspring. This might be related to loss of Rex1, but could also be the result of a broader role for RLIM in embryonic development [9]. Here, we explored the developmental consequences of RLIM deficiency in male embryos and feti and show that this protein is important for late pulmonary development and probably also for heart development.

Lung development is divided into five overlapping stages which are tightly regulated by a concerted action of different signaling pathways including BMP pathway [27,28]. Among the numerous BMP ligands, BMP4 has been identified as a major lung development regulator in mice [20,27,28,29,30]. We showed that *Rlim* deletion causes a retarded branching morphogenesis already present at an early stage in two out of four E16.5 feti, with impaired terminal sacculation, a delayed surfactant production and a subsequent defective lung inflation. The pulmonary vascular development seems normal in KO mice, indicating that the observed paleness is most likely due to an increased pulmonary vascular resistance in collapsed lungs.

By targeting SMAD7 for degradation, RLIM has been demonstrated to increase TGFβ, BMP and activin signaling activity in mouse embryonic stem cells, mouse embryonic fibroblasts and in zebrafish embryos where SMAD7 is suggested to be a specific RLIM substrate [5]. RLIM mediated promotion of TGFβ signaling has also been demonstrated in U2OS cell line via the direct interaction between RLIM and the SMURF2 ubiquitin ligase which interferes with SMURF2-SMAD7 interaction that otherwise causes degradation of the TGFβ receptors [4]. In addition, SMURF2 interferes with R-SMADs including SMAD1, SMAD2 and SMAD3 [31]. However, RLIM biological function in mammalian organogenesis has never been explored. Using lung lysates, we addressed RLIM involvement in BMP signaling and confirmed the positive correlation between endogenous RLIM level and BMP activity as indicated by an increase of SMAD7 and phosphorylated SMAD1/5/9 depletion in RLIM deficient lungs.

The lung phenotype resulting from *Rlim* knockout resembles the phenotypes that have previously been reported in studies exploring BMP pathway involvement in lung development. Using lung explant culture, Chen et al have demonstrated the negative impact of knocking down SMAD1 on lung branching morphogenesis, epithelial cells proliferation and differentiation [32]. Interestingly, these same anomalies have been observed when SMURF1 is overexpressed in lung epithelium, which was accompanied with SMAD1 and SMAD5 reduction [33]. SMURF1 is another SMAD specific E3 Ubiquitin Ligase that inhibits BMP signaling. In mammalian cells, SMURF1 expression induces the ubiquitination and proteasome-mediated degradation of SMAD1 and SMAD5 in a ligand independent manner [34] [35]. Moreover, SMURF1 controls the intracellular signaling of BMPs through interaction with SMAD6 and SMAD7 and induces BMP type I receptor, SMAD1 and SMAD5 degradation [36]. Whilst SMAD5 was decreased in KO lungs, SMAD1 showed unchanged expression. By analogy with the established RLIM-SMURF2 interaction [4], we hypothesize that RLIM does not only activate BMP pathway through SMAD7 degradation, but also through interaction with SMURF1 (Fig 9). SMAD5 reduction could result from a decreased RLIM-SMURF1 interaction and therefore an enhanced SMURF1 activity, either induced or not by accumulated SMAD7. SMAD1 insensitivity to RLIM depletion (and to the suggested resulting SMURF1 upregulation) could be explained either by a requirement of a threshold of the endogenous SMAD1 level to activate SMURF1 mediated degradation or by another layer of a yet to be uncovered regulation.

**Fig 9:**
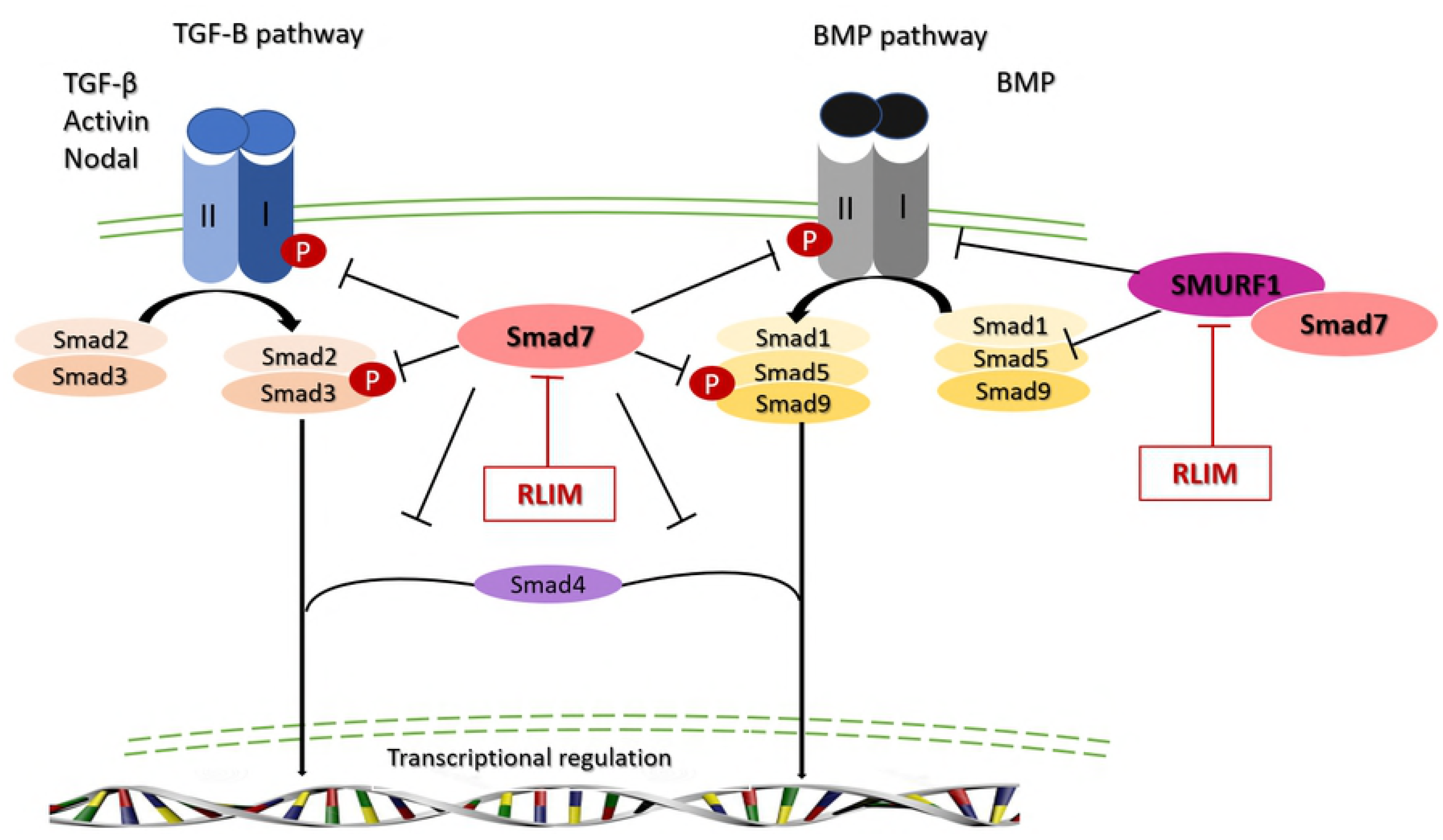
Suggested model for positive regulation of BMP pathway by RLIM. RLIM targets the inhibitory SMAD7 for degradation and probably interacts with SMURF1 which targets BMP type I receptor, SMAD1 and SMAD5 for degradation.

Based on the other previously reported *Rlim* mouse models, which have the same genetic background and approximately the same deletion size [7,10], male lethality is most likely due to *Rlim* deletion in visceral endoderm. Rather than being completely displaced by the epiblast-derived definitive endoderm during gastrulation, some visceral endoderm cells have been demonstrated to remain dispersed by the intercalating definitive endoderm cells. These visceral endoderm cells proliferate to become included in the embryonic gut epithelium [11], and ultimately contribute to gut-derived tissues including liver, pancreas and brain [38]. In addition to these tissues, lung epithelium is also derived from the foregut endoderm whereas lungs mesenchyme is a mesodermal structure [28] which does not involve visceral endoderm derived cell. Hence, although RLIM, SMAD1 and SMURF1 are expressed in both epithelial and mesenchymal cells [32,33,39], we expect the phenotype of our mouse model to only result from an abnormal BMP4 signaling in epithelial cells. However, a mesenchymal anomaly cannot be excluded since lung development requires an intimate epithelial-mesenchymal cells crosstalk [40,41].

To elucidate the mechanism by which BMP4 regulates lung development, Xu et al have used epithelial-specific Smad1 and SMAD5 KO mice [20]. Importantly, epithelial-specific Smad5 KO did not result in any abnormality suggesting that SMAD5 reduction in our lung samples is not underlying the observed anomalies. On the other hand, the study of epithelial-specific *Smad1* KO mouse model has shown a downregulation of its target gene WIF1[20]. Despite the dramatic reduction of phosphorylated SMAD1/5/9, WIF1 and ID3 expression was comparable in KO and WT lungs, suggesting a SMAD independent regulation of this gene.

Fifty percent of the spontaneously born *Rnf12* KO males survived, indicating an incomplete penetrance of the observed pulmonary phenotype. This incomplete penetrance is probably due to a genetic background effect, which is well documented in TGF-β field [42]. In support of this hypothesis, mortality rate of CAST/Ei mice carrying the same *Rnf12* deletion is higher than that of C57BL/6 mice [9]. On the other hand, the survival rate of KO male pups in our study largely exceeds that of the Cesarian section dissected E18.5 KO males of which 16 out of 17 deceased. In addition to the abnormal lung maturation, the growth restriction observed in the Rnf12^-/y^ KO mice may have contributed to an aggravation of the respiratory insufficiency in the pre-term dissected pups [43]. It is of note that the only mutant survivor had the highest birth weight of all, around 1.4 g, suggesting a critical birth weight of perinatal survival. Fetal growth restriction can be due to a placental abnormality [44]. Placentas of E16.5 KO mice showed a normal gross morphology. Specifically, placenta size and shape as well as umbilical cord vessel insertion was comparable in all E16.5 feti (data not shown).

Except for growth restriction and possibly heart defects, this *Rnf12* mouse model does not phenocopy the human phenotype. However, our study provides new clues about the molecular mechanism behind *RLIM* lesions associated human congenital malformations. In two of the reported nine families, lung hypoplasia was described in association with congenital diaphragmatic hernia [14]. Lung specimen of these patients were not examined. In general, lungs of patients with CDH are immature with thickened alveolar walls, and epithelial differentiation failure, an increased interstitial tissue due to defective apoptosis and compromised vascular growth [45]. These defects are more severe in CDH ipsilateral lung. The TGFβ pathway is also a key player in lung branching and morphogenesis [46] and has been the focus of the very few studies on human CDH related lung pathology. Similarly to normal human developing lung, SMAD2/3 mediated TGFβ signaling increases throughout pregnancy in CDH associated hypoplastic lungs [47]. *RLIM* pathogenic variants in these families may have reduced the canonical SMAD2/3 signaling in lungs with a subsequent dysregulation of downstream genes or pathways and possibly additional lung anomalies.

In conclusion, RLIM is important for lung development in mice and exerts this function in part through positive regulation of the intracellular propagation of BMP signaling. The murine phenotype does not phenocopy the human phenotype and may thus not be the best animal model to understand *RLIM* pathology.

## Material and methods

### Mouse strain

Mouse procedures were approved by the ethical committee and performed according to the guidelines of the Animal Welfare Committee of KU Leuven, Belgium (P248/2015).

C57BL/6 conventional *Rlim* KO mice used in this study and genotyping have previously been described [9]. All examined mice and embryos were derived from natural matings. For genotyping, DNA was extracted from ear biopsies of postnatal pups and from yolk sac or tail biopsies of collected embryos and feti. Genotypes were determined by PCR and gel electrophoresis using the previously described primers [48]. Pregnant KO females were euthanized by cervical dislocation at E9.5, E16.5 or E18.5. Unless specified otherwise, feti where euthanized by decapitation.

E9.5 and E16.5 embryos were imaged, sampled for genotyping and processed for subsequent analysis. A subset of E16.5 feti was subjected to microdissection to search for gross malformations. E18.5 feti were recovered by caesarian section and monitored for 1 hour while being warmed on a 37°C heating plate. Color, presence of heart beat, breathing rate, movements, reactivity to foot pinch and weight were recorded. The surviving neonates were euthanized after 1 hour and lung samples were collected for subsequent analysis.

To compare E18.5 prenatal lung morphology without air exposure, some litters were euthanized in-utero by immersion of the uterus in ice cold PBS [18] and fixed. Lung samples were subsequently histopathologically examined.

### Histological procedures

E16.5 embryos and E18.5 lungs samples were fixed in 4% paraformaldehyde in PBS overnight at 4°C, dehydrated, paraffin embedded and sectioned (5 μm). Sections were used for hematoxylin-eosin (H&E), periodic acid-Schiff (PAS) and immunohistochemistry staining. PAS staining was performed by the UZ Leuven Pathology department.

Immunohistochemistry staining was performed using rat anti-CD31 (Pharmingen 553370) and rat anti-endomucin (EMCN) (SC-65495) according to standard procedures.

Microscopic images were acquired using Leica DM5500 microscope and processed with ImageJ software. Air space area, PAS positive and endomucin positive areas were measured by averaging a minimum of 10 views per fetus using a 20x objective for the former measurement and a 40x objective for the two latter ones.

### Antibodies and immunodetection

Western blot analysis was performed according to standard procedures. The primary antibodies were rabbit anti-RLIM (ABE1949), rabbi, rabbit anti-phospho-SMAD5 (ab76296), rabbit anti-SMAD1 (CST 9743), rabbit anti-SMAD5 (ab40771), rabbit anti-SMAD7 (PA1-41506), rabbit anti-WIF1 (ab155101) and rabbit anti-ID3 (sc-490). Chemiluminescence (ECL Reagent Plus or Ultra from Perkin Elmer) was detected using a BioRad imager. Band intensities were assessed after background extraction using ImageJ software and subsequently normalized to the corresponding Actin or Tubulin values.

### Statistics

R package was used for statistical calculations. Binomial test (Fig 1) and Mann-Whitney U test (Figs 3, 5 and 6) were used to determine statistical significance. A p-value below 0.05 was considered statistically significant.

## Acknowledgement

The authors wish to thank Mrs Greet Peeters and Mrs Annick Francis for their technical support. The Leica DM5500 microscope was provided by InfraMouse (KU Leuven-VIB) through a Hercules type 3 project (ZW09-03).

This work was made possible by grants from the KU Leuven (SymBioSys (PFV/10/016), GOA/12/015 to J.R.V and C12/16/023 to A.Z), the Hercules foundation (ZW11-14) and from the Belgian Science Policy OfficeInteruniversity Attraction Poles (BELSPO-IAP) programme through the project IAP P7/43-BeMGI to J.R.V. MK is supported by the Erasmus+Program of the European Union (Framework agreement number 2013-0040). This publication reflects the views only of the authors, and the European Commission cannot be held responsible for any use that may be made of the information contained therein.

## References

1. Bach I, Rodriguez-Esteban C, Carrière C, Bhushan A, Krones A, Rose DW, et al. RLIM inhibits functional activity of LIM homeodomain transcription factors via recruitment of the histone deacetylase complex. Nat Genet. 1999;22(4):394–9.

2. Her YR, Chung IK. Ubiquitin ligase RLIM modulates telomere length homeostasis through a proteolysis of TRF1. J Biol Chem. 2009;284(13):8557–66.

3. Gao K, Wang C, Jin X, Xiao J, Zhang E, Yang X, et al. RNF12 promotes p53-dependent cell growth suppression and apoptosis by targeting MDM2 for destruction. Cancer Lett. 2016;375(1):133–41.

4. Huang Y, Yang Y, Gao R, Yang X, Yan X, Wang C, et al. RLIM interacts with Smurf2 and promotes TGF-β induced U2OS cell migration. Biochem Biophys Res Commun. 2011;414(1):181–5.

5. Zhang L, Huang H, Zhou F, Schimmel J, Pardo CG, Zhang T, et al. RNF12 Controls Embryonic Stem Cell Fate and Morphogenesis in Zebrafish Embryos by Targeting Smad7 for Degradation. Mol Cell. 2012;46(5):650–61.

6. Jiao B, Ma H, Shokhirev M, Drung A, Yang Q, Shin J, et al. Paternal RLIM/Rnf12 is a survival factor for milk-producing alveolar cells. Cell. 2012;149(3):630–41.

7. Shin J, Bossenz M, Chung Y, Ma H, Byron M, Taniguchi-Ishigaki N, et al. Maternal Rnf12/RLIM is required for imprinted X-chromosome inactivation in mice. Nature. 2010;467(7318):977–81.

8. Gontan C, Achame EM, Demmers J, Barakat TS, Rentmeester E, Van Ijcken W, et al. RNF12 initiates X-chromosome inactivation by targeting REX1 for degradation. Nature. 2012;485(7398):386–90.

9. Gontan C, Mira-Bontenbal H, Magaraki A, Dupont C, Barakat TS, Rentmeester E, et al. REX1 is the critical target of RNF12 in imprinted X chromosome inactivation in mice. Nat Commun. 2018;9(1):4752.

10. Shin J, Wallingford MC, Gallant J, Marcho C, Jiao B, Byron M, et al. RLIM is dispensable for X-chromosome inactivation in the mouse embryonic epiblast. Nature. 2014;511(7507):86–9.

11. Kwon GS, Viotti M, Hadjantonakis AK. The Endoderm of the Mouse Embryo Arises by Dynamic Widespread Intercalation of Embryonic and Extraembryonic Lineages. Dev Cell. 2008;15(4):509–20.

12. Tønne E, Holdhus R, Stansberg C, Stray-Pedersen A, Petersen K, Brunner HG, et al. Syndromic X-linked intellectual disability segregating with a missense variant in RLIM. Eur J Hum Genet. 2015;23(12):1652–6.

13. Hu H, Haas SA, Chelly J, Van Esch H, Raynaud M, De Brouwer APM, et al. X-exome sequencing of 405 unresolved families identifies seven novel intellectual disability genes. Mol Psychiatry. 2016;21(1):133–48.

14. Frints SGM, Ozanturk A, Criado GR, Grasshoff U, Hoon B, Field M, et al. Pathogenic variants in E3 ubiquitin ligase RLIM/RNF12 lead to a syndromic X-linked intellectual disability and behavior disorder. Mol Psychiatry. 2018;(Mim 300379).

15. Souchelnytskyi S, Nakayama T, Nakao A, Moren A, Heldin CH, Christian JL, et al. Physical and Functional Interaction of Murine andXenopus Smad7 with Bone Morphogenetic Protein Receptors and Transforming Growth Factor-Receptors. J Biol Chem. 1998;273(39):25364–70.

16. Dunwoodie SL, Rodriguez T a, Beddington RS. Msg1 and Mrg1, founding members of a gene family, show distinct patterns of gene expression during mouse embryogenesis. Mech Dev. 1998;72(1-2):27–40.

17. Gavrilov S, Lacy E. Genetic dissection of ventral folding morphogenesis in mouse: Embryonic visceral endoderm-supplied BMP2 positions head and heart. Curr Opin Genet Dev. 2013;23(4):461–9.

18. Jakus Z, Gleghorn JP, Enis DR, Sen A, Chia S, Liu X, et al. Lymphatic function is required prenatally for lung inflation at birth. J Exp Med. 2014;211(5):815–26.

19. Liu W, Selever J, Wang D, Lu M-F, Moses KA, Schwartz RJ, et al. Bmp4 signaling is required for outflow-tract septation and branchial-arch artery remodeling. Proc Natl Acad Sci. 2004;101(13):4489–94.

20. Xu B, Chen C, Chen H, Zheng S-G, Bringas P, Xu M, et al. Smad1 and its target gene Wif1 coordinate BMP and Wnt signaling activities to regulate fetal lung development. Development. 2011;138(5):925–35.

21. Reis LM, Tyler RC, Schilter KF, Abdul-Rahman O, Innis JW, Kozel BA, et al. BMP4 loss-of-function mutations in developmental eye disorders including SHORT syndrome. Hum Genet. 2011;130(4):495–504.

22. Jay PY, Bielinska M, Erlich JM, Mannisto S, Pu WT, Heikinheimo M, et al. Impaired mesenchymal cell function in Gata4 mutant mice leads to diaphragmatic hernias and primary lung defects. Dev Biol. 2007;301(2):602–14.

23. Miyazaki Y, Oshima K, Fogo A, Ichikawa I. Evidence that bone morphogenetic protein 4 has multiple biological functions during kidney and urinary tract development. Kidney Int. 2003;63(3):835–44.

24. Reis GS Dos, Simões E Silva AC, Freitas IS, Heilbuth TR, Marco LA De, Oliveira EA, et al. Study of the association between the BMP4 gene and congenital anomalies of the kidney and urinary tract. J Pediatr (Rio J). 2014;90(1):58–64.

25. Chen T, Li Q, Xu J, Ding K, Wang Y, Wang W, et al. Mutation screening of BMP4, BMP7, HOXA4 and HOXB6 genes in Chinese patients with hypospadias. Eur J Hum Genet. 2007;15(1):23–8.

26. Wang H, Zhang L, Wang N, Zhu H, Han B, Sun F, et al. Next-generation sequencing reveals genetic landscape in 46, XY disorders of sexual development patients with variable phenotypes. Hum Genet. 2018;137(3):265–77.

27. Herriges M, Morrisey EE. Lung development: orchestrating the generation and regeneration of a complex organ. Development. 2014;141(3):502–13.

28. Chao C-M, El Agha E, Tiozzo C, Minoo P, Bellusci S. A Breath of Fresh Air on the Mesenchyme: Impact of Impaired Mesenchymal Development on the Pathogenesis of Bronchopulmonary Dysplasia. Front Med. 2015;2:1–13.

29. Alejandre-Alcázar MA, Shalamanov PD, Amarie O V., Sevilla-Pérez J, Seeger W, Eickelberg O, et al. Temporal and spatial regulation of bone morphogenetic protein signaling in late lung development. Dev Dyn. 2007;236(10):2825–35.

30. Ameis D, Khoshgoo N, Keijzer R. Abnormal lung development in congenital diaphragmatic hernia. Semin Pediatr Surg. 2017;26(3):123–8.

31. Lin X, Liang M, Feng XH. Smurf2 is a ubiquitin E3 ligase mediating proteasome-dependent degradation of Smad2 in transforming growth factor-β signaling. J Biol Chem. 2000;275(47):36818–22.

32. Chen C, Chen H, Sun J, Bringas P, Chen Y, Warburton D, et al. Smad1 expression and function during mouse embryonic lung branching morphogenesis. Am J Physiol Cell Mol Physiol. 2005;288(6):L1033–9.

33. Shi W, Chen H, Sun J, Chen C, Zhao J, Wang Y-L, et al. Overexpression of Smurf1 negatively regulates mouse embryonic lung branching morphogenesis by specifically reducing Smad1 and Smad5 proteins. Am J Physiol Cell Mol Physiol. 2004;286(2):L293–300.

34. Zhu H, Kavsak P, Abdollah S, Wrana JL, Thomsen GH. A SMAD ubiquitin ligase targets the BMP pathway and affects embryonic pattern formation. Nature. 1999;400(6745):687–93.

35. Wrana JL, Attisano L. The Smad pathway. Cytokine Growth Factor Rev. 2000;11(1-2):5–13.

36. Murakami G, Watabe T, Takaoka K, Miyazono K, Imamura T. Cooperative Inhibition of Bone Morphogenetic Protein Signaling by Smurf1 and Inhibitory Smads. Mol Biol Cell. 2003;14:2809–2817.

37. Shin J, Wallingford MC, Gallant J, Marcho C, Jiao B, Byron M, et al. RLIM is dispensable for X-chromosome inactivation in the mouse embryonic epiblast. Nature. 2014;511(7507):86–9.

38. Viotti M, Nowotschin S, Hadjantonakis AK. Afp::mCherry, a red fluorescent transgenic reporter of the mouse visceral endoderm. Genesis. 2011;49(3):124–33.

39. Du Y, Guo M, Whitsett JA, Xu Y. “LungGENS”: A web-based tool for mapping single-cell gene expression in the developing lung. Thorax. 2015;70(11):1092–4.

40. Yamamoto H, Jun Yun E, Gerber HP, Ferrara N, Whitsett JA, Vu TH. Epithelial-vascular cross talk mediated by VEGF-A and HGF signaling directs primary septae formation during distal lung morphogenesis. Dev Biol. 2007;308(1):44–53.

41. Hines EA, Sun X. Tissue crosstalk in lung development. J Cell Biochem. 2014;115(9):1469–77.

42. Bonyadi M, Rusholme SA, Cousins FM, Su HC, Biron CA, Farrall M, et al. Mapping of a major genetic modifier of embryonic lethality in TGF b1 knockout mice. Nat Genet. 1997;15(2):207–11.

43. Turgeon B, Meloche S. Interpreting Neonatal Lethal Phenotypes in Mouse Mutants: Insights Into Gene Function and Human Diseases. Physiol Rev. 2009;89(1):1–26.

44. Ward JM, Elmore SA, Foley JF. Pathology methods for the evaluation of embryonic and perinatal developmental defects and lethality in genetically engineered mice. Vet Pathol. 2012;49(1):71–84.

45. Donahoe PK, Longoni M, High FA. Polygenic Causes of Congenital Diaphragmatic Hernia Produce Common Lung Pathologies. Am J Pathol. 2016;186(10):2532–43.

46. Bartram U, Speer CP. The Role of Transforming Growth Factor β in Lung Development and Disease. Chest. 2004;125(2):754–65.

47. Vuckovic A, Herber-Jonat S, Flemmer AW, Ruehl IM, Votino C, Segers V, et al. Increased TGF-β: a drawback of tracheal occlusion in human and experimental congenital diaphragmatic hernia? Am J Physiol - Lung Cell Mol Physiol. 2016;310(4):L311–27.

48. Jonkers I, Barakat TS, Achame EM, Monkhorst K, Kenter A, Rentmeester E, et al. RNF12 Is an X-Encoded Dose-Dependent Activator of X Chromosome Inactivation. Cell. 2009;139(5):999–1011.

